# Activation of the pro-resolving receptors Fpr2/3 attenuate lipopolysaccharide-induced inflammatory microglial activation

**DOI:** 10.1101/2020.07.09.195198

**Authors:** Edward S. Wickstead, Bradley T. Elliott, Christopher Biggs, Stephen J. Getting, Simon McArthur

## Abstract

Neuroinflammation driven primarily by microglia directly contributes to neuronal death in many neurodegenerative diseases. Classical anti-inflammatory approaches aim to suppress pro-inflammatory mediator production, but exploitation of inflammatory resolution, the active endogenous process whereby inflammation is terminated, may also be of benefit. A key driver of peripheral inflammatory resolution, formyl peptide receptors 2/3 (Fpr2/3), is expressed by microglia, but its therapeutic potential in neurodegeneration remains unclear. Here, we sought to provide a proof-of-principle that Fpr2/3 targeting could reverse inflammatory microglial activation induced by the potent bacterial inflammogen lipopolysaccharide (LPS). Stimulation of murine BV2 microglia with LPS triggered release of nitric oxide, TNFα and IL-10, upregulated surface expression of CD38 and CD40 and suppressed CD206, all of which were reduced by subsequent treatment with the Fpr2/3 ligand C43. Cellular exposure to LPS also stimulated NADPH oxidase and mitochondrial derived reactive oxygen species production, effects reversed by C43 treatment. Mechanistic studies showed C43 to act through p38 MAPK phosphorylation and reduction of LPS-induced NFκB nuclear translocation through prevention of IκBα degradation. Here, we provide proof-of-concept data indicating exploitation of the pro-resolving receptor Fpr2/3 as a promising therapeutic strategy in neuroinflammatory conditions.

## Introduction

Neuroinflammation is a complex biological response required to protect the central nervous system (CNS) from injury or infection. Microglia, the resident macrophage-like innate immune cells of the CNS, are essential to this process (Bellver-Landete et al., 2019; Heckmann et al., 2019), with shifts in cellular activation being crucial to eliminate the noxious insult, but also to facilitate inflammatory resolution and tissue repair (Zabala et al., 2018). However, if this homeostatic resolution response becomes impaired, it can result in the eventual development of chronic neuroinflammation. This has been reported following trauma, ischaemic-reperfusion injury or during neurodegenerative disease (Chamorro et al., 2016; Gyoneva and Ransohoff, 2015; Heneka et al., 2015).

Resolution, the process through which the acute inflammatory response is curtailed and tissue homeostasis restored, is an endogenous part of the immune response, driven to a large degree by pro-inflammatory mediators (Serhan, 2017). Although most widely studied in the peripheral immune response, it is thought that similar circuitry underlies the microglial response in neuroinflammation (Dokalis and Prinz, 2019). Thus, actively stimulating inflammatory resolution rather than simply inhibiting pro-inflammatory mediators could hold promise as a new treatment approach for the neuroinflammatory pathology associated with many neurodegenerative diseases (Frigerio et al., 2018).

The formyl peptide receptors (FPRs) are G-protein coupled receptors with roles in chemotaxis, host defence and inflammation (Dalli et al., 2012; McArthur et al., 2015). Of these, FPR2 and its murine functional homologues Fpr2/3 have a key role in peripheral inflammatory resolution (Dufton et al., 2010; Gobbetti et al., 2014; Stempel et al., 2016), with central functions in the regulation of monocyte/macrophage recruitment (Drechsler et al., 2015; Gobbetti et al., 2014; McArthur et al., 2015), phenotype and behaviour (Chen et al., 2018; McArthur et al., 2020; Scannell et al., 2007). Important protective actions have been identified in diverse inflammatory settings, including viral infection (Schloer et al., 2019), mucosal injury (Birkl et al., 2019) and acute heart failure (Kain et al., 2019). The receptor is expressed in numerous immune cells, including microglia (Yu and Ye, 2014), but the role it might play in neuroinflammation has been relatively poorly examined. Previous work has shown the major anti-inflammatory ligand for FPR2, the protein annexin A1 (ANXA1) to attenuate neuroinflammation in murine haemorrhagic (Ding et al., 2019) or ischaemic (Liu et al., 2015) stroke models, to promote microglial *β*-amyloid removal (Ries et al., 2016) and to associate with inflammation and disease severity in individuals with multiple sclerosis (Colamatteo et al., 2019) and Alzheimer’s disease (Thygesen et al., 2019).

Given this evidence for beneficial effects of FPR2 ligands in neuroinflammatory disease, we sought to investigate the relationship between the receptor and microglial behaviour, in a model of inflammatory activation, *in vitro* exposure to bacterial lipopolysaccharide (LPS). In particular, we investigated whether treatment of microglia after inflammatory stimulation could reverse their activation and promote a return to homeostasis, a key feature of any potential therapeutic agent in the treatment of neuroinflammatory disease.

## Materials & Methods

### Drugs and reagents

The FPR2 agonist Compound-43 (*N*-(4-Chlorophenyl)-*N*-[2,3-dihydro-1-methyl-5-(1-methylethyl)-3-oxo-2-phenyl-1*H*-pyrazol-4-yl]-urea) and antagonist WRW_4_ (Trp-Arg-Trp-Trp-Trp-NH_2_) were purchased from Tocris Ltd, UK. Isolated and purified lipopolysaccharides from *Escherichia coli*, serotype O111:B4 were purchased from Merck Millipore Ltd, UK.

### Cell culture

The BV2 murine microglial line and shRNA annexin A1 clones were kind gifts from Prof. E. Blasi (Università degli Studi di Modena e Reggio Emilia, Italy) and Dr. Egle Solito (The William Harvey Research Institute, Queen Mary University of London, UK), respectively. Cells were cultured in DMEM supplemented with 5% heat-inactivated fetal calf serum (FCS) with 100 μM non-essential amino acids, 2 mM L-alanyl-L-glutamine and 50 mg/ml penicillin-streptomycin (Thermofisher Scientific, UK) at 37°C with 5% CO_2_ and 95% air. Prior to experimentation, BV2 cells were serum starved for 24 h.

### Griess assay

Nitric oxide production was indirectly detected through the measurement of nitrite using the Griess method (Misko et al., 1993). Cellular supernatant was treated with Griess reagent (3.85 μM napthylethylenediamine dihydrochloride, 58.1 μM sulphanilamide, 5% ortho-phosphoric acid) at room temperature in the dark for 15 min. Absorbance was detected at 540 nm with a CLARIOstar microplate reader (BMG Labtech, Germany). Final concentrations were determined by reference to a standard curve of sodium nitrite in cell culture medium (3-100 μM).

### Cytokine ELISA

Supernatant content of tumour necrosis factor alpha (TNFα) and interleukin-10 (IL-10) were assayed by murine-specific sandwich ELISAs using commercially available kits, according to the manufacturer’s instructions (ThermoFisher Scientific, UK). A CLARIOstar spectrophotometer (BMG Labtech, Germany) was used to measure absorbance at 450 nm. Absorbance values recorded at 570 nm were subtracted to reduce optical interference.

### p38 MAPK ELISA

BV2 cells were treated according to experimental design. Samples were lysed and protein normalised following content assessment by the Bradford reagent (Walker and Kruger, 2003) for use in an InstantOne™ total/phosphor multispecies p38 ELISA kit (ThermoFisher Scientific, UK) according to the manufacturer’s instructions. Absorbance was measured at 450 nm and 570 nm, as previously described.

### Phenotypic marker expression

Following incubation with blocking buffer (0.01 M PBS,1 μg/ml anti rat anti-mouse CD16/32 monoclonal antibody, 1% FCS, 1mM CaCl_2_) on ice for 15 min, BV2 cells were labelled with APC-conjugated rat anti-mouse CD38 (1:80) and CD40 (1:20), and PE-conjugated rat anti-mouse CD206 (1:40), and FITC-conjugated rat anti-mouse CD86 (1:80; Biolegend, UK) for 30 min on ice for analysis by flow cytometry. Immunofluorescence was analysed for 10,000 singlet events per sample using a BD FACSCanto II (BD Biosciences, UK) flow cytometer; data were analysed using FlowJo 8.8.1 software (Treestar Inc., CA, USA).

### E. coli bioparticle phagocytosis

Microglial phagocytic capacity was determined using BODIPY FL conjugated *Escherichia coli* (K-12 strain) bioparticles (ThermoFisher Scientific, UK). Following experimental treatments, cells were incubated with bioparticle conjugates at a ratio of 50 particles per cell in PBS for 30 min at 37°C in the dark. Cells were washed with 4°C PBS and fluorescence of non-engulfed particles was quenched with the addition of 0.2% Trypan blue (ThermoFisher Scientific, UK) for 1 min. Cells were washed and collected, and cellular fluorescence was determined using a FACSCanto II flow cytometer (BD Biosciences, UK) equipped with a 488nm laser and FlowJo 8.8.1 software (Treestar Inc. CA, USA). A total of 10,000 single events were quantified per sample.

### Lactate & glucose determination

L-lactate production and glucose usage were simultaneously determined using a YSI 2300 Stat Plus machine (YSI Life Sciences Inc, UK). Following experimental treatments, cell supernatant was collected and stored at -80°C until required. Concentration readings for L-lactate and glucose were linear up to 30 mM/L and 50 mM/L, respectively.

### Reactive oxygen species (ROS) assays

Total intracellular ROS production was quantified using 6-chloromethyl-2’,7’-dichlorodihydrofluorescein diacetate, acetyl ester (CM-H_2_DCFDA; ThermoFisher Scientific, UK) according to the manufacturer’s instructions. Cells were plated at 200,000 cells/cm^2^ in phenol red-free (PRF) DMEM and serum starved overnight prior to being pre-loaded with 5 µM CM-H_2_DCFDA for 15 min at 37°C. Unbound dye was then removed, and fresh PRF-DMEM was added prior to experimentation. Following treatment administration, cellular fluorescence was determined every 5 min for 1 h at 37°C using a CLARIOstar fluorescence microplate reader (BMG Labtech, Germany) with excitation and emission filters set to 492nm and 517nm, respectively.

Mitochondrial superoxide production (mtROS) was quantified using the MitoSOX Red tracer (ThermoFisher Scientific, UK) according to the manufacturer’s instructions, with a loading concentration of 2.5 μM. Following treatment, cellular fluorescence was determined every 5 min for 1 h at 37°C using a CLARIOstar fluorescence microplate reader (BMG Labtech, Germany) with excitation and emission filters set at 510nm and 580nm, respectively.

### Western blot analysis

Samples boiled in 6x Laemmli buffer were subjected to standard SDS-PAGE (10%) prior to being electrophoretically blotted onto Immobilon-P polyvinylidene difluoride membranes (Merck, UK). Ponceau S staining (Merck, UK) was used to quantify total protein. Membranes were blotted using antibodies raised against murine superoxide dismutase 2 (SOD; rabbit monoclonal, 1:1000), haem oxygenase-1 (HO-1; rabbit polyclonal, 1:1000) or IκBα (mouse monoclonal, 1:1000; Cell Signaling Technology, Leiden, The Netherlands) in Tris-buffer saline solution containing 0.1% Tween-20 and 5% (w/v) non-fat dry milk overnight at 4°C. Membranes were subsequentially washed with Tris-buffer saline solution containing 0.1% Tween-20, prior to incubation with appropriate secondary antibody (horseradish peroxidase-conjugated goat anti-rabbit, 1:5000, ThermoFisher Scientific, UK), for 90 min at room temperature. Proteins were then detected using enhanced chemiluminescence detection (0.4 mM p-coumaric acid, 7.56 mM H_2_O_2_ and 2.5 mM luminol in 1 M Tris, pH 8.5) and visualised on X-ray film (Scientific Laboratory Supplies Limited, Nottingham, UK).

### Immunofluorescence & confocal microscopy

Following experimental treatments, BV2 microglia in chambered culture slides were fixed by incubation with 2% formaldehyde in PBS for 10 min at 4°C, washed and non-specific antibody binding was minimised through 30 min incubation at room temperature in PBS containing 10% hiFCS and 0.05% Triton-X 100 (ThermoFisher Scientific, UK). Cells were then incubated with rabbit anti-mouse p67phox monoclonal antibody (1:500, clone EPR5064, Abcam Ltd, UK) and mouse anti-mouse gp91phox monoclonal antibody (1:50, clone 53, BD Biosciences, UK) or rabbit anti-mouse NFκB monoclonal antibody (1:400, clone D14E12, Cell Signaling Technologies, Leiden, The Netherlands) overnight at 4°C in PBS with 1% hiFCS and 0.05% Triton-X 100. Cells were washed and incubated with AF488-conjugated goat anti-mouse and AF647-conjugated goat anti-rabbit secondary antibodies (both 1:500, ThermoFisher Scientific, UK) in PBS with 1% hiFCS and 0.05% Triton-X 100 at room temperature in the dark for 1 h. Cells were washed with PBS, nuclei stained with 180 nM DAPI in ddH_2_O for 5 min and mounted with Mowiol solution. Cells were imaged using an LSM710 confocal microscope (Leica, UK) fitted with 405nm, 488nm, and 647nm lasers and a 63x oil immersion objective lens. Images were captured with ZEN Black software (Zeiss, UK) prior to analysis with ImageJ 1.51w (National Institutes of Health, USA). Co-localisation of p67phox and gp91phox was determined using graded lookup tables for red and green pixels only.

### Statistical analysis

Sample sizes were calculated to detect differences of 15% or more, with a power of 0.85 and α set at 5%; calculations being informed by previously published data (Loiola et al., 2019; McArthur et al., 2010). All experimental data are presented as mean ± SEM, with a minimum of *n* = 3 independent cultures. All assays were performed in triplicate. For all data, the normality of distribution was established with the Shapiro-Wilke test, followed by further analysis with two-tailed Student’s t tests to compare two groups or, for multiple comparison analysis, one-, two-, or three-way ANOVA followed by Tukey’s HSD *post hoc* test. A p value of <0.05 was considered statistically significant. All statistical analyses were performed with Graph Pad Prism 8 software (GraphPad Software, CA, USA).

## Results

### Fpr2/3 reverses the pro-inflammatory actions of LPS in BV2 microglia

The potent pro-inflammatory stimulus LPS was selected as a model inflammogen, at a concentration (50 ng/ml) based upon our previous research (Loiola et al., 2019; McArthur et al., 2010). Administration of the selective Fpr2/3 agonist C43 (100 nM) 1 h after LPS treatment reversed LPS-induced production of nitrite and TNFα at both 24 and 48 h (Figure 1A-B), although without affecting the LPS-induced increase in iNOS expression (Suppl. Figure 1). A similar pro-resolving effect was seen on assessment of IL-10 production, with C43 potentiating its release compared to LPS alone at 48 h post-treatment (Figure 1C). Moreover, these pro-resolving effects of C43 were sensitive to 10 min pre-treatment with the Fpr2/3 specific antagonist WRW_4_ at 10 µM (Figure 1D-1F), further confirming the involvement of this receptor.

**Figure 1:**
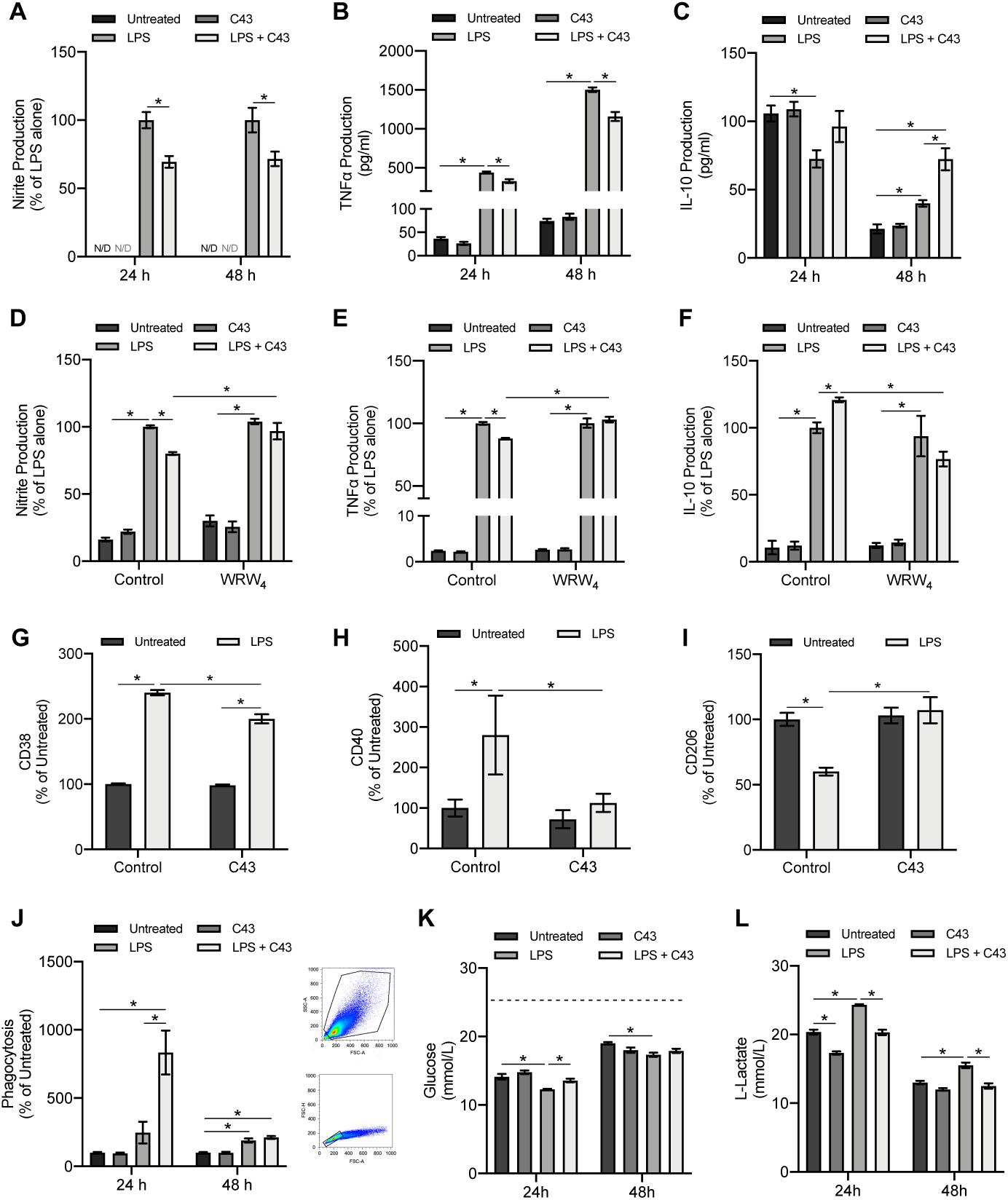
LPS induced microglial inflammation is reduced by C43 through switching to a pro-resolving phenotype. **A, B**) LPS (50 ng/ml) increases the release of the pro-inflammatory mediators nitric oxide and TNFα in microglia after 24 and 48 h exposure. 1 h post-treatment with C43 (100 nM) significantly reduced these increases. **C**) C43 elicits the increase in IL-10 production at 48 h following LPS administration when compared to untreated and LPS treated cells alone. **D-F**) the effects of C43 were blocked by 10 min pre-treatment with the selective Fpr2/3 antagonist, WRW_4_ (10 µM). Administration of LPS upregulates the expression of the pro-inflammatory phenotypic markers (**G**) CD38 and (**H**) CD40, which reversed by administration of C43 24 h afterwards. **I**) LPS reduced the pro-resolving marker CD206, which was also reversed by C43 treatment. **J**) C43 post-treatment increases *E. coli* fluorescent bioparticle phagocytosis by upwards of 8-fold compared to untreated but has no effect without initial LPS induction. Cell gating strategies to remove microglial doublets are shown. **K**) LPS increases BV2 glucose utilisation after 24 h and 48 h exposure, as represented by the reduction in supernatant glucose content. 1 h post-treatment with C43 significantly reversed this at 24 h only. Dashed line represents glucose concentration in fresh DMEM. **L**) LPS increased cellular L-lactate production at 24 and 48 h, but 1 h post-treatment with C43 significantly reversed this back to baseline levels at both time points. Data are mean ± SEM of 3-7 independent cultures in triplicate; * *P*<0.05.

As endogenously produced ANXA1 is a critical pro-resolving mediator for numerous immune cell types and is expressed by microglia (Loiola et al., 2019; McArthur et al., 2010), we examined whether the effects of C43 were mediated through this protein. However, no changes in ANXA1 expression were identified, and shRNA for ANXA1 had no effect on the cytokine changes elicited by C43 (Suppl. Figure 2).

**Figure 2:**
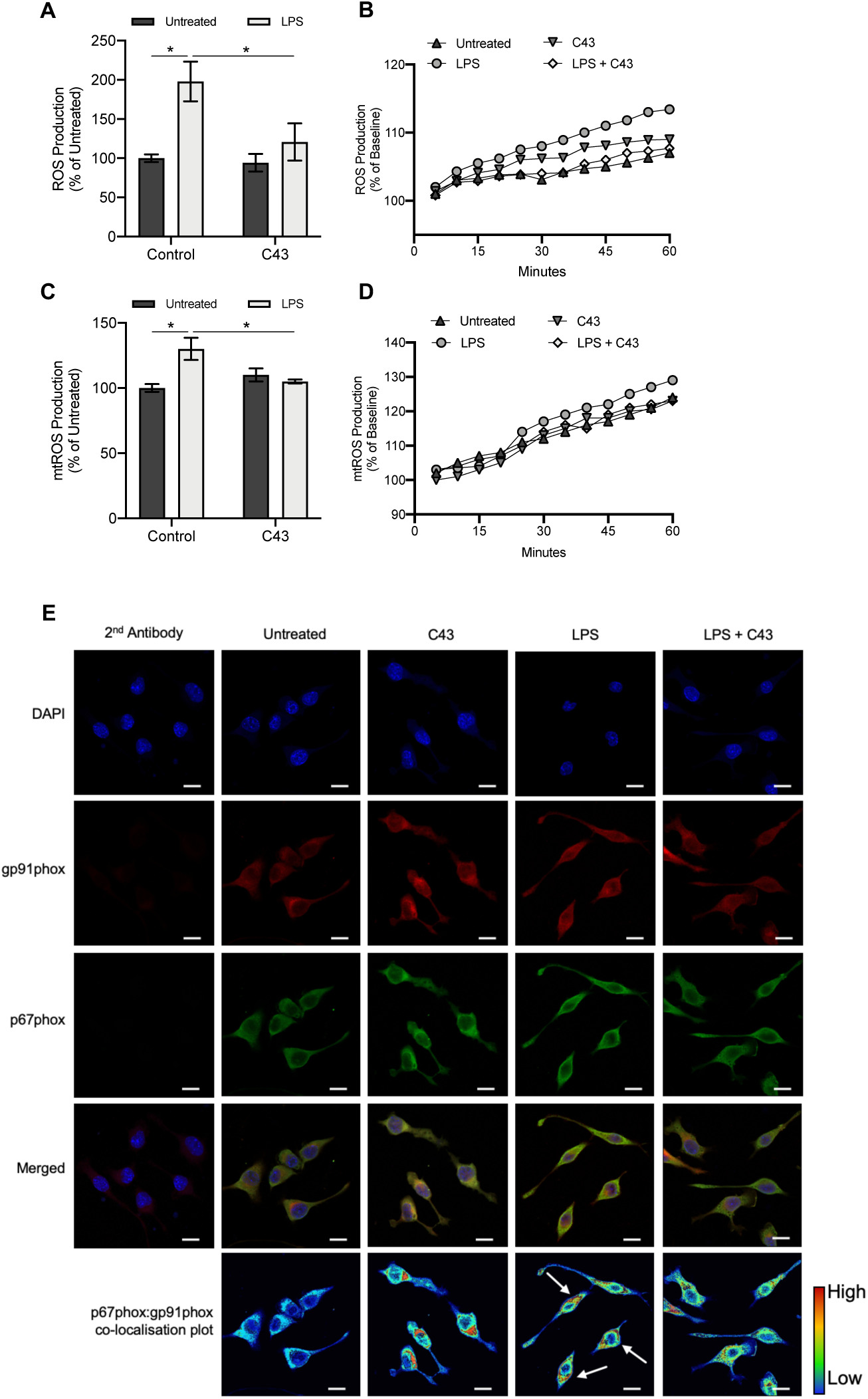
Fpr2/3 reverses LPS-induced ROS production through modulation of the mitochondria and NADPH oxidase. **A, B**) addition of LPS (50 ng/ml) for 1 h increases ROS production in BV2 microglia but is completely reversed by 10 min post-treatment with C43 (100 nM). **C, D**) mitochondrial ROS (mtROS) production elicited by LPS exposure for 1 h was also successfully reversed by C43 treatment. **E**) treatment of BV2 cells for 30 min with 50 ng/ml LPS stimulated the plasma membrane co-localisation of the NADPH oxidase subunits p67phox (green) and gp91phox (red), an effect reversed by treatment with 100 nM C43 administered 10 min post-LPS. Nuclei are counterstained with DAPI (blue). Co-localisation intensity of p67phox and gp91phox is represented by the false-colour plots. Graphical data are means ± SEM of 3-5 independent cultures, plated in triplicate. Confocal images are representative of 3 independent cultures; scale bar = 10 μm; * *P*<0.05.

The expression of different cell surface markers provide insight into the activation state of microglia, hence we examined whether C43 treatment could modify expression of pro-inflammatory (CD38, CD40, CD86) and pro-resolving (CD206) phenotypic markers when given 24 h post-LPS administration. Although CD86 did not respond to LPS stimulation (data not shown), expression of both CD38 and CD40 were upregulated by LPS (Figure 1G-H), with CD38 and CD40 being reversed 24h after subsequent C43 treatment (Figure 1G-H). Surface expression of the anti-inflammatory, pro-resolution marker CD206 was suppressed following LPS treatment, but this was rescued by subsequent C43 treatment (Figure 1I). C43 had no effect on any of these markers when administered alone.

An important aspect of inflammation is the removal of cellular debris and invading micro-organisms by phagocytosis, a process we and others have previously shown to involve Fpr2/3 in peripheral immune cells (Gobbetti et al., 2014; Wan et al., 2014; Weiß et al., 2019). We therefore investigated whether this would similarly apply to microglia. Treatment with C43 alone did not affect BV2 cell capacity to phagocytose BODIPY FL-conjugated *E. coli* bioparticles, but the Fpr2/3 agonist when given 1 h after administration of LPS significantly increased bacterial uptake approximately 8-fold after 24 h (Figure 1J), confirming that activation of the receptor can stimulate processes aiming to terminate inflammation.

The role of metabolic pathways in governing immune cell function has steadily become apparent over recent years, with inflammatory cells tending to favour glycolysis over oxidative phosphorylation for energy generation (Kelly et al., 2015; O’Neill and Pearce, 2016). We therefore investigated whether Fpr2/3 activation could modulate the effects of LPS stimulation upon glucose utilisation and L-lactate production, an indirect measure of glycolysis (Brand et al., 2016). LPS significantly increased glucose utilisation in BV2 microglia, whereas C43 administration 1 h post-LPS reversed this at 24 h only (Figure 1K). Addition of C43 also reversed the LPS-induced increase in L-lactate at both 24 and 48 h, with C43 treatment alone reducing L-lactate production at 24 h compared to untreated (Figure 1L). Taken together, these data strongly indicate that selective Fpr2/3 activation with C43 can significantly reverse and diminish LPS-elicited microglial activation.

### LPS induces ROS production through both the mitochondria and NADPH oxidase activation, a response reversed by Fpr2/3 agonist C43

A key anti-bacterial property of immune cell activation is the production of reactive oxygen species (ROS) in response to stimulation by bacterial components such as LPS (Kim et al., 2010). However, uncontrolled ROS production can be highly damaging to bystander neurons and can be a significant cause of neuroinflammatory damage (Cobb and Cole, 2015). To determine whether Fpr2/3 activation could influence LPS-induced ROS production, BV2 microglia were pre-treated with LPS for 10 min, followed by addition of C43 and cellular ROS production was monitored every 5 min for 1 h. LPS increased ROS production by approximately 100% compared to untreated, with C43 completely reversing this (Figure 2A-B). Approximately 30% of the observed increase in ROS following LPS stimulation could be accounted for by mitochondrial ROS production, with this also being reversed by C43 administration (Figure 2C-D).

Aside from mitochondrial production, the main source of microglial ROS in response to inflammatory stimulation is through activation of NADPH oxidase, also termed NOX2 (Huang et al., 2018). NOX2 is a multi-subunit enzyme, wherein activation requires the translocation of the p67phox subunit from the cytosol to the plasma membrane-bound gp91phox subunit (Haslund-Vinding et al., 2017). Confocal microscopic analysis of BV2 cells stimulated with LPS for 30 min indicated clear co-localisation of p67phox and gp91phox subunits at the plasma membrane, an effect that was prevented by treatment with C43 10 min post-LPS (Figure 2E). These Fpr2/3 mediated effects were independent of the antioxidant systems involving HO-1 and SOD2, with no changes in their expression (Suppl. Figure 3). Thus, selective Fpr2/3 stimulation can successfully reverse LPS-induced ROS production via both mitochondrial and NADPH oxidase associated mechanisms.

**Figure 3:**
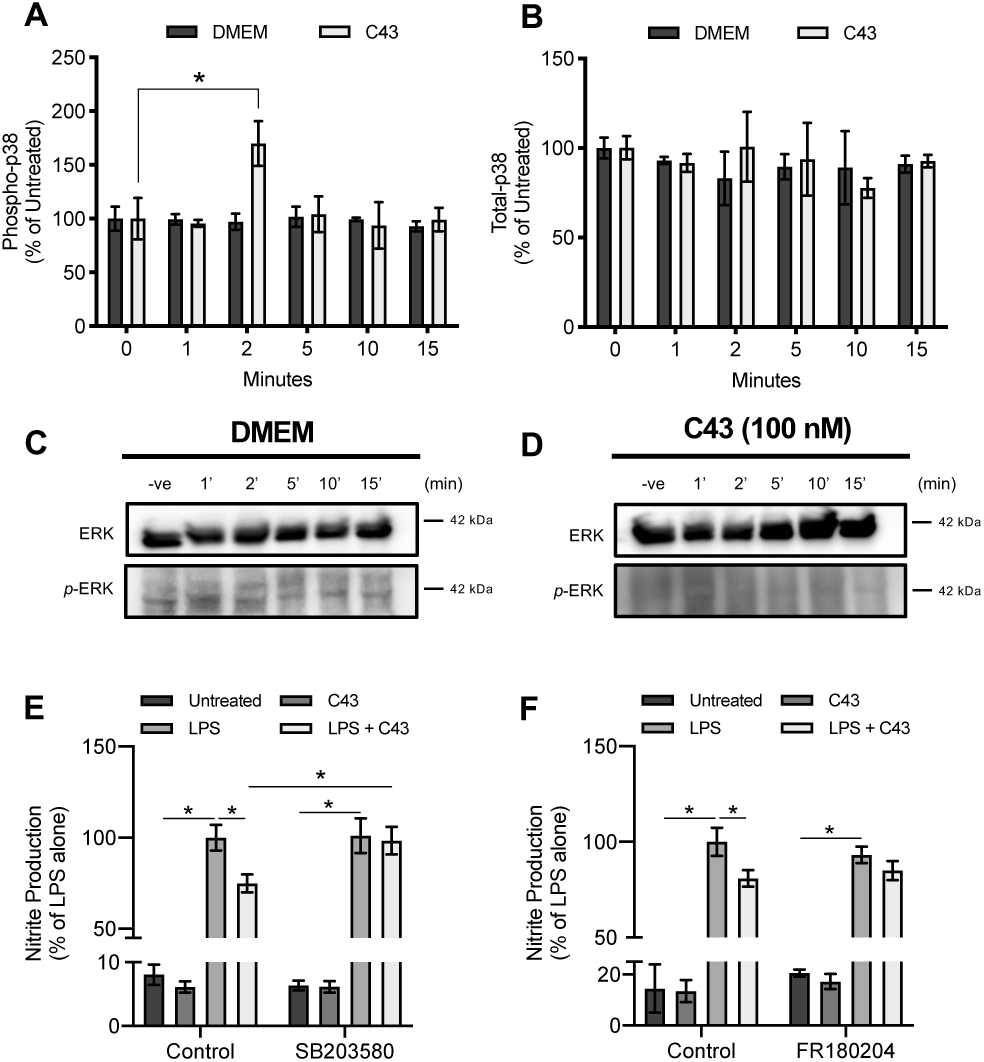
Fpr2/3 binding with C43 facilitates signalling through p38 MAPK, but not ERK1/2. **A, B)** addition of C43 (100 nM) for 2 min increased the phosphorylation of p38 MAPK by over 150% when compared to untreated. Cells administered with serum-free DMEM were used as a control. **C, D)** no changes were seen for ERK1/2 phosphorylation over the same administration period of C43. **E, F**) confirming the importance of p38 MAPK in facilitating the anti-inflammatory effects of C43, pre-treatment with the selective p38 inhibitor SB203580 (2 µM) but not the ERK1/2 inhibitor FR180204 (10 µM) 10 min prior to C43 prevents the nitrite reducing effects of this Fpr2/3 agonist; * *P*<0.05.

### C43 signals through p38 MAPK but not ERK1/2

We have previously identified that the Fpr2/3 agonist ANXA1 can upregulate IL-10 production via a p38 MAPK dependent mechanism (Cooray et al., 2013). To determine whether this was true for C43, p38 MAPK phosphorylation in BV2 lysates was determined by ELISA. Confirming this idea, 2 min treatment with C43 significantly increased p38 MAPK phosphorylation (Figure 3A & 3B). However, Fpr2/3 can also signal through ERK1/2 (Cooray et al., 2013; Kim et al., 2019), thus we also examined ERK1/2 phosphorylation. However, compared to p38, C43 did not activate ERK1/2 over the same time period (Figure 3C & 3D). To further corroborate this, the Griess assay was repeated at 48 h with both LPS and C43 as described previously, but with the inclusion of the selective p38 inhibitor SB203580 (2 µM) or the ERK1/2 inhibitor FR180204 (10 µM) 10 min prior to C43 treatment. Notably, SB203580 but not FR180204, inhibited the nitrite reducing effects of C43 (Figure 3E & 3F), further validating a role for p38 MAPK in mediating the anti-inflammatory effects of C43.

### C43 reduces NFκB activation by decreasing LPS-induced IκBα degradation

In microglia, LPS can trigger the production of many pro-inflammatory mediators including NO and TNFα, through the nuclear translocation of NFκB (Cao et al., 2018; Youssef et al., 2019). Confocal analysis of BV2 cells following LPS exposure confirmed nuclear translocation of the NFκB p65 subunit, an effect reversed by treatment with C43 10 minutes post-LPS (Figure 4A). This effect of C43 was inhibited by inclusion of the p38 inhibitor SB203580 (2 µM) (Figure 4A). NFκB p65 is held in the cytoplasm by the regulatory IκBα complex (Liu et al., 2017), proteolytic degradation of which can be stimulated by LPS-induced phosphorylation (Aslanidis et al., 2015). Analysis confirmed an LPS-induced reduction of cellular IκBα expression, which was prevented by treatment with C43 10 minutes post-LPS (Figure 4B-C), indicating that the anti-inflammatory effects of C43 are mediated through p38 MAPK activation and inhibition of NFκB nuclear translocation.

**Figure 4:**
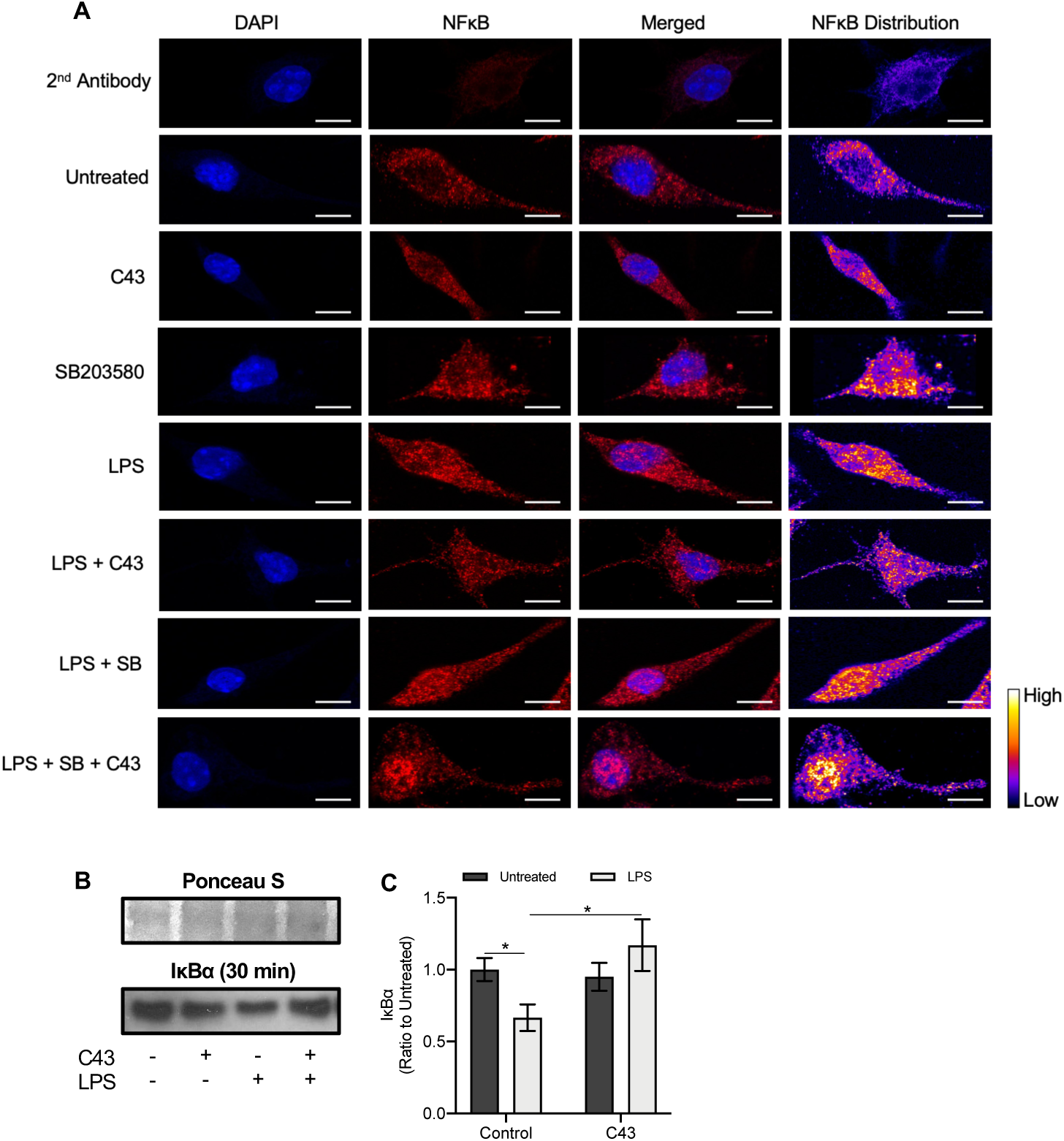
Fpr2/3 partially reverses LPS-induced NFκB nuclear translocation by preventing IκBα degradation. **A)** 30 min exposure of BV2 microglia to 50 ng/ml LPS stimulated the nuclear translocation of the p65 subunit of NFκB (red), an effect partially reversed by 10 min post-treatment of 100 nM C43. These observations of C43 were prevented with the inclusion of the p38 inhibitor SB203580 (2 µM) 5 min before. Nuclei are counterstained with DAPI (blue). **B, C**) 30 min exposure with 50 ng/ml LPS triggers NFκB p65 nuclear translocation through the degradation of IκBα, an effect prevented by the 10 min post-treatment of 100 nM C43. Confocal images are representative of 3 independent cultures, Scale bar = 10 µm. Graphical data are means ± SEM of 3-4 independent cultures, plated in triplicate; * *P*<0.05.

## Discussion

Alongside their central roles in neural circuit development, synaptic pruning and plasticity (Bar and Barak, 2019; Paolicelli et al., 2011), microglia are critical for tissue maintenance, the response to injury (Gyoneva and Ransohoff, 2015) and defence against invading pathogens (Ransohoff and El Khoury, 2015). However, hyperactivation of microglia can become pathological, leading to chronic neuroinflammation and neurodegeneration (Hickman et al., 2018). Evidence for the role of neuroinflammation in brain disease is widespread (Salter and Stevens, 2017), with clinical reports emphasising the significance of neuroinflammation in neurodegenerative conditions such as stroke (Meng et al., 2019), multiple sclerosis (Schattling et al., 2019), and Alzheimer’s disease (Felsky et al., 2019). This is also underlined through improved disease outcomes following the application of anti-inflammatory and immunomodulatory therapeutic approaches (Moussa et al., 2017; Smith et al., 2018; Tankou et al., 2018). Due to the potential for microglial activation to become damaging, strategies to control excessive inflammatory actions through promotion of a pro-resolving phenotype may be of significant benefit for a wide range of neurological diseases. In this study, we have used an *in vitro* cellular model to provide proof-of-principle evidence that targeting the pro-resolving receptor Fpr2/3 is a viable approach to restraining inflammatory microglial behaviour.

Murine Fpr2/3 and human FPR2 have central roles in the resolution of peripheral inflammation (Cooray et al., 2013; Gobbetti et al., 2014; McArthur et al., 2020, 2015; Vital et al., 2016). Despite also being expressed in microglia (Chen et al., 2007; McArthur et al., 2010; Ries et al., 2016), little is known about the receptor’s potential as an anti-neuroinflammatory target. Here we provide further evidence for the pro-resolving effects of Fpr2/3 agonists (Corminboeuf and Leroy, 2015), and extend these to CNS immune cell populations. Notably, our finding that post-LPS treatment with an Fpr2/3 specific agonist can significantly attenuate and reverse induction of a pro-inflammatory phenotype in BV2 microglia suggests that this receptor may have potential for exploitation as a therapeutic, rather than a prophylactic target. In particular, we suggest that activation of Fpr2/3 has potential to protect tissue from ongoing neuroinflammatory damage and is therefore a valuable target for further investigation of *in vivo* models of neuroinflammatory disease.

Our data moreover, indicate an important role for Fpr2/3 in restraining NFκB signalling via inhibition of IκBα degradation, extending our understanding of the receptor’s intracellular signalling pathways (Cooray et al., 2013), as well as highlighting its potential to modulate pro-inflammatory signalling. The modulation of LPS-induced NFκB by Fpr2/3 stimulation has been reported recently in both monocyte (de Gaetano et al., 2019) and osteoclast (Hu et al., 2020) cell lines, suggesting it may be a common feature of macrophage-lineage cells. As microglia are known to rapidly upregulate Fpr2/3 expression following inflammatory insult (Kirkley et al., 2017), this serves as a further indication for the importance of this receptor in inflammatory regulation, emphasising the concept that inflammatory resolution is an intrinsic, programmed part of the inflammatory response (Serhan and Savill, 2005).

In addition, we report for the first time that Fpr2/3 activation can reverse LPS-induced ROS production from two independent sources: the mitochondria and NADPH oxidase. Previous studies report the ability of LPS to induce ROS production in microglia via these pathways (Yauger et al., 2019; Zhang et al., 2014), both of which may be crucial in shifting microglial metabolic parameters (Nair et al., 2019) and triggering neuroinflammation (Hou et al., 2019; Park et al., 2015). This may be central to microglial associated neurodegeneration (Cheng et al., 2018; Ye et al., 2017). Moreover, in addition to their clear role in neurodegeneration (Hickman et al., 2018), microglia may well become pathological during other chronic inflammatory conditions. For example, microglia have been shown to be crucial in the induction of chronic pain in mice in the absence of nerve injury (Zhou et al., 2019). Thus, modulating microglia phenotype may hold additional premise in disorders wherein neurons do not become inherently damaged. Future work will determine whether the *in vitro* findings reported here can be translated into *in vivo* models. If this holds true, Fpr2/3 agonists capable of dampening pathological microglial activity could hold therapeutic promise for numerous neurological diseases.

## Conclusions

This study has identified that the Fpr2/3 agonist C43 is a potent pro-resolving ligand, successfully modulating microglial phenotype and inflammatory cytokine release alongside ablating mitochondrial and NADPH oxidase induced ROS production, following an LPS-induced inflammatory response. This data therefore suggests that the modulation of Fpr2/3 could hold potential in the development of therapeutics for neuroinflammatory disease.

## List of abbreviations

FPR2: Human formyl peptide receptor 2
Fpr2/3: Murine formyl peptide receptors 2/3
IL-10: Interleukin 10
LPS: Lipopolysaccharide
ROS: Reactive oxygen species
TNFα: Tumour necrosis factor alpha
WRW4: Trp-Arg-Trp-Trp-Trp-Trp

## Acknowledgements

EW was supported by a PhD Scholarship from the University of Westminster.

## Figure Legends

**Supplementary Figure 1.**
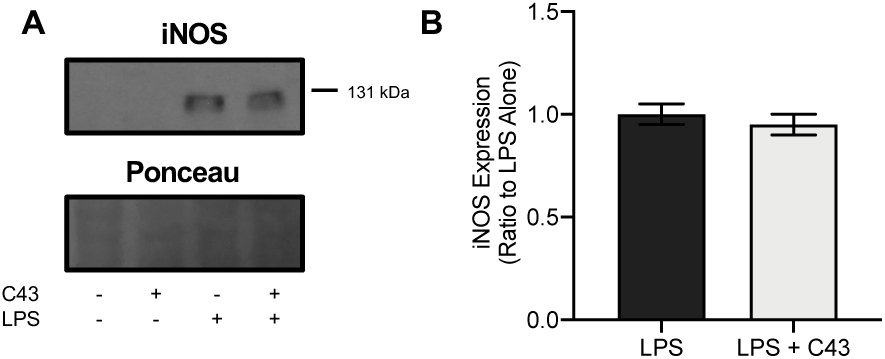
Reduction of nitrite by C43 is independent of iNOS regulation. **A, B)** administration of 50 ng/ml LPS for 24 h upregulated the translation of iNOS compared to untreated, whereas 100 nM C43 had no effect on iNOS expression when administered either alone, or 1 h post-LPS. Data are means ± SEM of 3 independent cultures.

**Supplementary Figure 2:**
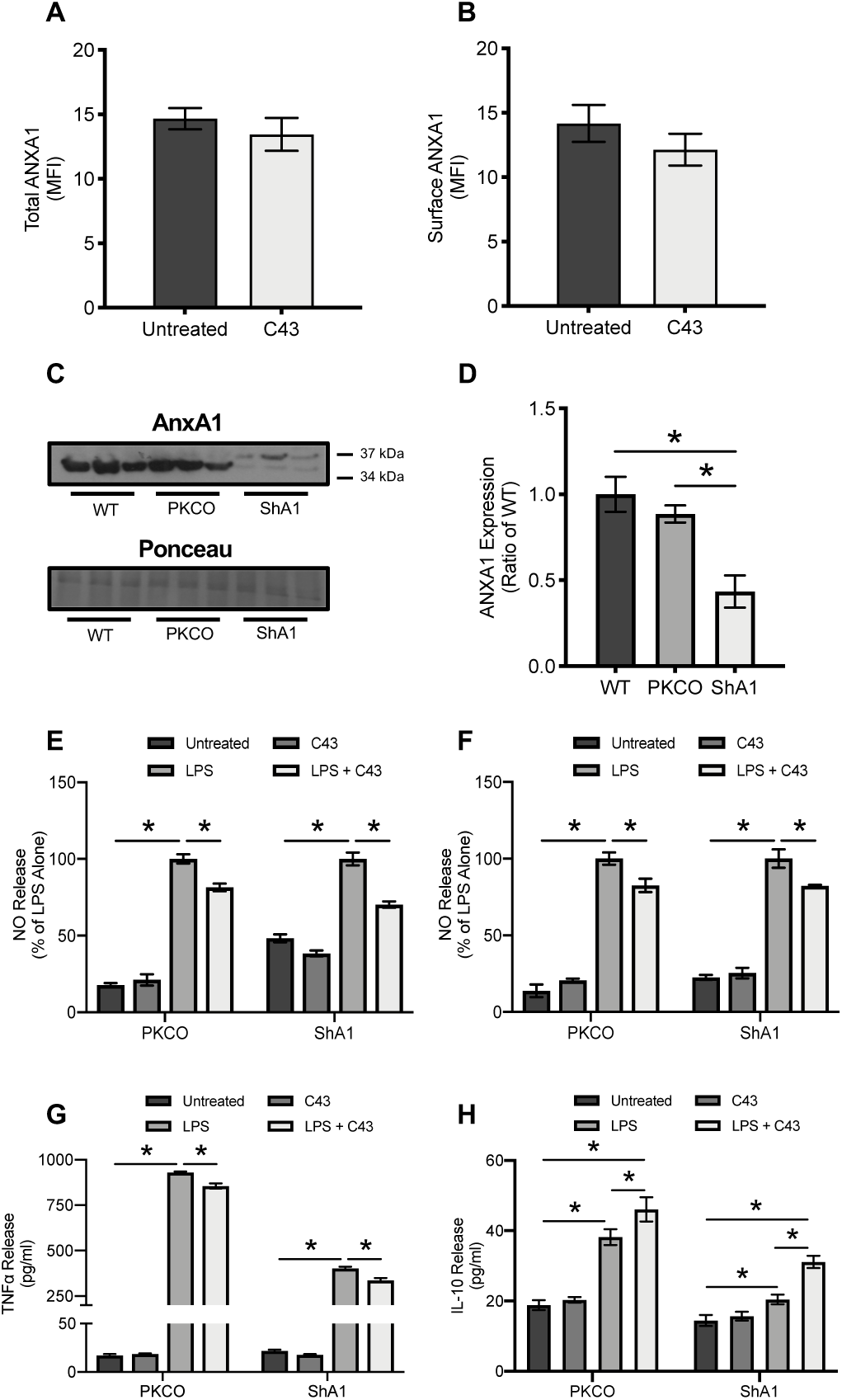
The anti-inflammatory effects of C43 are independent of ANXA1 regulation. **A, B**) the relative expression levels of both total and surface annexin A1 after 24 h treatment with 100 nM C43, determined with flow cytometry and 0.1% TX-100 cell permeabilization. **C, D**) transfection with ShA1 successfully reduced annexin A1 expression in BV2 microglia compared to untransfected and control plasmid transfection (PKCO). **E, F**; the anti-inflammatory nitrite reducing effects of 1 h post-treatment of C43 were retained even in PKCO and ShA1 transfected cells at 24 and 48 h. **G, H**; the effects of C43 on TNFα and IL-10 release at 48 h post-LPS administration were also retained in plasmid transfected microglia. Data are means ± SEM of 3-6 independent cultures, plated in triplicate; * *P*<0.05.

**Supplementary Figure 3:**
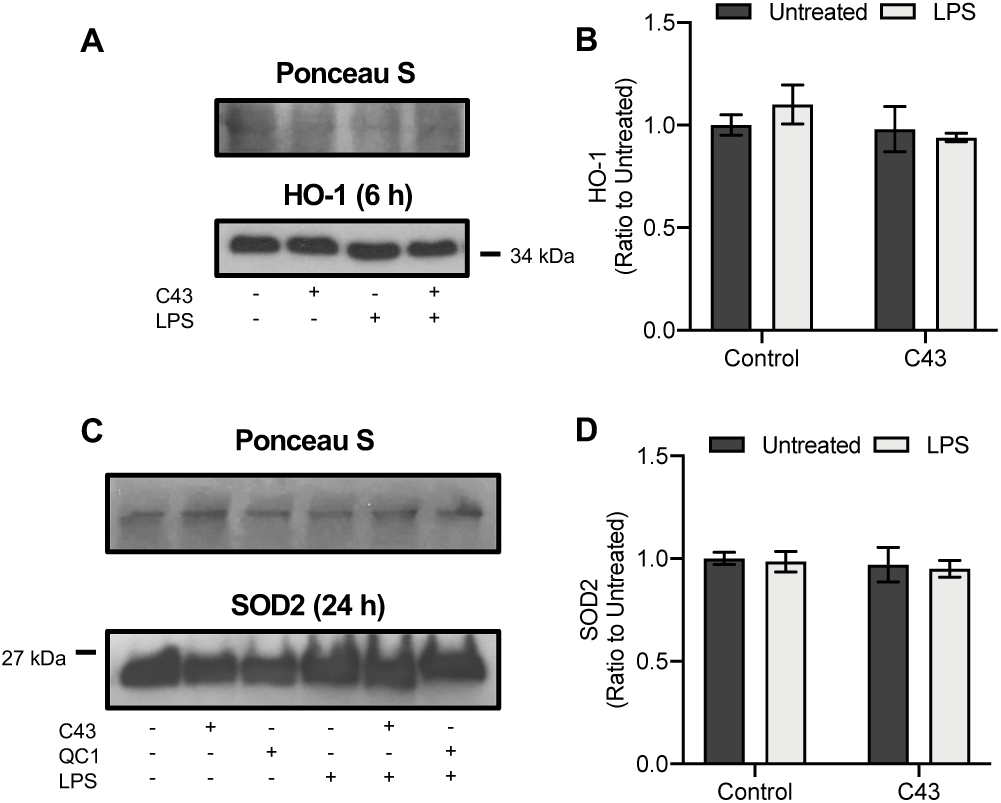
Neither LPS nor C43 treatment affected two major cellular antioxidant enzymes. **A, B**) no difference in HO-1 expression after 6 h exposure to LPS was observed in BV2 microglia, with and without the 1 h post treatment of C43. **C, D**) SOD2 expression remained unchanged after 24 h LPS addition, with C43 treatment also unable to modulate protein expression. Western blot analysis is representative of 3 independent cultures, with samples normalized to Ponceau S-defined total protein content; densitometric analysis data are means ± SEM of 3 independent cultures.

## References

Aslanidis, A., Karlstetter, M., Scholz, R., Fauser, S., Neumann, H., Fried, C., Pietsch, M., Langmann, T., 2015. Activated microglia/macrophage whey acidic protein (AMWAP) inhibits NFκB signaling and induces a neuroprotective phenotype in microglia. J. Neuroinflammation 12, 77. doi:10.1186/s12974-015-0296-6

Bar, E., Barak, B., 2019. Microglia roles in synaptic plasticity and myelination in homeostatic conditions and neurodevelopmental disorders. Glia 67, 2125–2141. doi:10.1002/glia.23637

Bellver-Landete, V., Bretheau, F., Mailhot, B., Vallières, N., Lessard, M., Janelle, M.-E., Vernoux, N., Tremblay, M.-È., Fuehrmann, T., Shoichet, M.S., Lacroix, S., 2019. Microglia are an essential component of the neuroprotective scar that forms after spinal cord injury. Nat. Commun. 10, 518. doi:10.1038/s41467-019-08446-0

Birkl, D., O’Leary, M.N., Quiros, M., Azcutia, V., Schaller, M., Reed, M., Nishio, H., Keeney, J., Neish, A.S., Lukacs, N.W., Parkos, C.A., Nusrat, A., 2019. Formyl peptide receptor 2 regulates monocyte recruitment to promote intestinal mucosal wound repair. FASEB J. 33, 13632–13643. doi:10.1096/fj.201901163R

Brand, A., Singer, K., Koehl, G.E., Kolitzus, M., Schoenhammer, G., Thiel, A., Matos, C., Bruss, C., Klobuch, S., Peter, K., Kastenberger, M., Bogdan, C., Schleicher, U., Mackensen, A., Ullrich, E., Fichtner-Feigl, S., Kesselring, R., Mack, M., Ritter, U., Schmid, M., Blank, C., Dettmer, K., Oefner, P.J., Hoffmann, P., Walenta, S., Geissler, E.K., Pouyssegur, J., Villunger, A., Steven, A., Seliger, B., Schreml, S., Haferkamp, S., Kohl, E., Karrer, S., Berneburg, M., Herr, W., Mueller-Klieser, W., Renner, K., Kreutz, M., 2016. LDHA-Associated Lactic Acid Production Blunts Tumor Immunosurveillance by T and NK Cells. Cell Metab. 24, 657–671. doi:10.1016/j.cmet.2016.08.011

Cao, X., Jin, Y., Zhang, H., Yu, L., Bao, X., Li, F., Xu, Y., 2018. The Anti-inflammatory Effects of 4-((5-Bromo-3-chloro-2-hydroxybenzyl) amino)-2-hydroxybenzoic Acid in Lipopolysaccharide-Activated Primary Microglial Cells. Inflammation 41, 530–540. doi:10.1007/s10753-017-0709-z

Chamorro, Á., Dirnagl, U., Urra, X., Planas, A.M., 2016. Neuroprotection in acute stroke: targeting excitotoxicity, oxidative and nitrosative stress, and inflammation. Lancet Neurol. 15, 869–881. doi:10.1016/S1474-4422(16)00114-9

Chen, K., Iribarren, P., Huang, J., Zhang, L., Gong, W., Cho, E.H., Lockett, S., Dunlop, N.M., Wang, J.M., 2007. Induction of the Formyl Peptide Receptor 2 in Microglia by IFN- and Synergy with CD40 Ligand. J. Immunol. 178, 1759–1766. doi:10.4049/jimmunol.178.3.1759

Chen, Y.-C., Lin, M.-C., Lee, C.-H., Liu, S.-F., Wang, C.-C., Fang, W.-F., Chao, T.-Y., Wu, C.-C., Wei, Y.-F., Chang, H.-C., Tsen, C.-C., Chen, H.-C., Taiwan Clinical Trial Consortium of Respiratory Disease (TCORE) group, 2018. Defective formyl peptide receptor 2/3 and annexin A1 expressions associated with M2a polarization of blood immune cells in patients with chronic obstructive pulmonary disease. J. Transl. Med. 16, 69. doi:10.1186/s12967-018-1435-5

Cheng, L., Chen, L., Wei, X., Wang, Y., Ren, Z., Zeng, S., Zhang, X., Wen, H., Gao, C., Liu, H., 2018. NOD2 promotes dopaminergic degeneration regulated by NADPH oxidase 2 in 6-hydroxydopamine model of Parkinson’s disease. J. Neuroinflammation 15, 243. doi:10.1186/s12974-018-1289-z

Cobb, C.A., Cole, M.P., 2015. Oxidative and nitrative stress in neurodegeneration. Neurobiol. Dis. 84, 4–21. doi:10.1016/J.NBD.2015.04.020

Colamatteo, A., Maggioli, E., Azevedo Loiola, R., Hamid Sheikh, M., Calì, G., Bruzzese, D., Maniscalco, G.T., Centonze, D., Buttari, F., Lanzillo, R., Perna, F., Zuccarelli, B., Mottola, M., Cassano, S., Galgani, M., Solito, E., De Rosa, V., 2019. Reduced Annexin A1 Expression Associates with Disease Severity and Inflammation in Multiple Sclerosis Patients. J. Immunol. ji1801683. doi:10.4049/jimmunol.1801683

Cooray, S.N., Gobbetti, T., Montero-Melendez, T., McArthur, S., Thompson, D., Clark, A.J.L., Flower, R.J., Perretti, M., 2013. Ligand-specific conformational change of the G-protein-coupled receptor ALX/FPR2 determines proresolving functional responses. Proc. Natl. Acad. Sci. U. S. A. 110, 18232–7. doi:10.1073/pnas.1308253110

Corminboeuf, O., Leroy, X., 2015. FPR2/ALXR Agonists and the Resolution of Inflammation. J. Med. Chem. 58, 537–559. doi:10.1021/jm501051x

Dalli, J., Montero-Melendez, T., McArthur, S., Perretti, M., 2012. Annexin A1 N-Terminal Derived Peptide Ac2-26 Exerts Chemokinetic Effects on Human Neutrophils. Front. Pharmacol. 3, 28. doi:10.3389/fphar.2012.00028

de Gaetano, M., Butler, E., Gahan, K., Zanetti, A., Marai, M., Chen, J., Cacace, A., Hams, E., Maingot, C., McLoughlin, A., Brennan, E., Leroy, X., Loscher, C.E., Fallon, P., Perretti, M., Godson, C., Guiry, P.J., 2019. Asymmetric synthesis and biological evaluation of imidazole- and oxazole-containing synthetic lipoxin A4 mimetics (sLXms). Eur. J. Med. Chem. 162, 80–108. doi:10.1016/j.ejmech.2018.10.049

Ding, Y., Flores, J., Klebe, D., Li, P., McBride, D.W., Tang, J., Zhang, J.H., 2019. Annexin A1 attenuates neuroinflammation through FPR2/p38/COX-2 pathway after intracerebral hemorrhage in male mice. . Neurosci. Res. jnr.24478. doi:10.1002/jnr.24478

Dokalis, N., Prinz, M., 2019. Resolution of neuroinflammation: mechanisms and potential therapeutic option. Semin. Immunopathol. 41, 699–709. doi:10.1007/s00281-019-00764-1

Drechsler, M., de Jong, R., Rossaint, J., Viola, J.R., Leoni, G., Wang, J.M., Grommes, J., Hinkel, R., Kupatt, C., Weber, C., Döring, Y., Zarbock, A., Soehnlein, O., 2015. Annexin A1 counteracts chemokine-induced arterial myeloid cell recruitment. Circ. Res. 116, 827–35. doi:10.1161/CIRCRESAHA.116.305825

Dufton, N., Hannon, R., Brancaleone, V., Dalli, J., Patel, H.B., Gray, M., D’Acquisto, F., Buckingham, J.C., Perretti, M., Flower, R.J., 2010. Anti-inflammatory role of the murine formyl-peptide receptor 2: ligand–specific effects on leukocyte responses and experimental inflammation. J. Immunol. 184, 2611–9. doi:10.4049/jimmunol.0903526

Felsky, D., Roostaei, T., Nho, K., Risacher, S.L., Bradshaw, E.M., Petyuk, V., Schneider, J.A., Saykin, A., Bennett, D.A., De Jager, P.L., 2019. Neuropathological correlates and genetic architecture of microglial activation in elderly human brain. Nat. Commun. 10, 409. doi:10.1038/s41467-018-08279-3

Frigerio, F., Pasqualini, G., Craparotta, I., Marchini, S., van Vliet, E.A., Foerch, P., Vandenplas, C., Leclercq, K., Aronica, E., Porcu, L., Pistorius, K., Colas, R.A., Hansen, T. V, Perretti, M., Kaminski, R.M., Dalli, J., Vezzani, A., 2018. n-3 Docosapentaenoic acid-derived protectin D1 promotes resolution of neuroinflammation and arrests epileptogenesis. Brain 141, 3130–3143. doi:10.1093/brain/awy247

Gobbetti, T., Coldewey, S.M., Chen, J., McArthur, S., le Faouder, P., Cenac, N., Flower, R.J., Thiemermann, C., Perretti, M., 2014. Nonredundant protective properties of FPR2/ALX in polymicrobial murine sepsis. Proc. Natl. Acad. Sci. U. S. A. 111, 18685–90. doi:10.1073/pnas.1410938111

Gyoneva, S., Ransohoff, R.M., 2015. Inflammatory reaction after traumatic brain injury: therapeutic potential of targeting cell–cell communication by chemokines. Trends Pharmacol. Sci. 36, 471–480. doi:10.1016/j.tips.2015.04.003

Haslund-Vinding, J., McBean, G., Jaquet, V., Vilhardt, F., 2017. NADPH oxidases in oxidant production by microglia: activating receptors, pharmacology and association with disease. Br. J. Pharmacol. 174, 1733–1749. doi:10.1111/bph.13425

Heckmann, B.L., Teubner, B.J.W., Tummers, B., Boada-Romero, E., Harris, L., Yang, M., Guy, C.S., Zakharenko, S.S., Green, D.R., 2019. LC3-Associated Endocytosis Facilitates β-Amyloid Clearance and Mitigates Neurodegeneration in Murine Alzheimer’s Disease. Cell 178, 536-551.e14. doi:10.1016/j.cell.2019.05.056

Heneka, M.T., Carson, M.J., Khoury, J. El, Landreth, G.E., Brosseron, F., Feinstein, D.L., Jacobs, A.H., Wyss-Coray, T., Vitorica, J., Ransohoff, R.M., Herrup, K., Frautschy, S.A., Finsen, B., Brown, G.C., Verkhratsky, A., Yamanaka, K., Koistinaho, J., Latz, E., Halle, A., Petzold, G.C., Town, T., Morgan, D., Shinohara, M.L., Perry, V.H., Holmes, C., Bazan, N.G., Brooks, D.J., Hunot, S., Joseph, B., Deigendesch, N., Garaschuk, O., Boddeke, E., Dinarello, C.A., Breitner, J.C., Cole, G.M., Golenbock, D.T., Kummer, M.P., 2015. Neuroinflammation in Alzheimer’s disease. Lancet Neurol. 14, 388–405. doi:10.1016/S1474-4422(15)70016-5

Hickman, S., Izzy, S., Sen, P., Morsett, L., El Khoury, J., 2018. Microglia in Neurodegeneration. Nat. Neurosci. 21, 1359–1369. doi:10.1038/s41593-018-0242-x

Hou, L., Sun, F., Huang, R., Sun, W., Zhang, D., Wang, Q., 2019. Inhibition of NADPH oxidase by apocynin prevents learning and memory deficits in a mouse Parkinson’s disease model. Redox Biol. 22, 101134. doi:10.1016/j.redox.2019.101134

Hu, J., Li, X., Chen, Y., Han, X., Li, L., Yang, Z., Duan, L., Lu, H., He, Q., 2020. The protective effect of WKYMVm peptide on inflammatory osteolysis through regulating NF-κB and CD9/gp130/STAT3 signalling pathway. J. Cell. Mol. Med. 24, 1893–1905. doi:10.1111/jcmm.14885

Huang, W.Y., Lin, S., Chen, H.Y., Chen, Y.P., Chen, T.Y., Hsu, K. Sen, Wu, H.M., 2018. NADPH oxidases as potential pharmacological targets against increased seizure susceptibility after systemic inflammation. J. Neuroinflammation 15, 140. doi:10.1186/s12974-018-1186-5

Kain, V., Jadapalli, J.K., Tourki, B., Halade, G. V, 2019. Inhibition of FPR2 impaired leukocytes recruitment and elicited non-resolving inflammation in acute heart failure. Pharmacol. Res. 146, 104295. doi:10.1016/j.phrs.2019.104295

Kelly, B., Aj, L., Neill, O.’, 2015. Metabolic reprogramming in macrophages and dendritic cells in innate immunity. Nat. Publ. Gr. 25. doi:10.1038/cr.2015.68

Kim, S.-Y., Lee, J.-G., Cho, W.-S., Cho, K.-H., Sakong, J., Kim, J.-R., Chin, B.-R., Baek, S.-H., 2010. Role of NADPH oxidase-2 in lipopolysaccharide-induced matrix metalloproteinase expression and cell migration. Immunol. Cell Biol. 88, 197–204. doi:10.1038/icb.2009.87

Kim, Y.E., Park, W.S., Ahn, S.Y., Sung, D.K., Sung, S.I., Kim, J.H., Chang, Y.S., 2019. WKYMVm hexapeptide, a strong formyl peptide receptor 2 agonist, attenuates hyperoxia-induced lung injuries in newborn mice. Sci. Rep. 9, 6815. doi:10.1038/s41598-019-43321-4

Kirkley, K.S., Popichak, K.A., Afzali, M.F., Legare, M.E., Tjalkens, R.B., 2017. Microglia amplify inflammatory activation of astrocytes in manganese neurotoxicity. J. Neuroinflammation 14, 99. doi:10.1186/s12974-017-0871-0

Liu, J.-H., Feng, D., Zhang, Y.-F., Shang, Y., Wu, Y., Li, X.-F., Pei, L., 2015. Chloral Hydrate Preconditioning Protects Against Ischemic Stroke via Upregulating Annexin A1. CNS Neurosci. Ther. 21, 718–726. doi:10.1111/cns.12435

Liu, T., Zhang, L., Joo, D., Sun, S.-C., 2017. NF-κB signaling in inflammation. Signal Transduct. Target. Ther. 2, 17023. doi:10.1038/sigtrans.2017.23

Loiola, R.A., Wickstead, E.S., Solito, E., McArthur, S., 2019. Estrogen Promotes Pro-resolving Microglial Behavior and Phagocytic Cell Clearance Through the Actions of Annexin A1. Front. Endocrinol. (Lausanne). 10, 420. doi:10.3389/fendo.2019.00420

McArthur, S., Cristante, E., Paterno, M., Christian, H., Roncaroli, F., Gillies, G.E., Solito, E., 2010. Annexin A1: a central player in the anti-inflammatory and neuroprotective role of microglia. J. Immunol. 185, 6317–28. doi:10.4049/jimmunol.1001095

McArthur, S., Gobbetti, T., Kusters, D.H.M., Reutelingsperger, C.P., Flower, R.J., Perretti, M., 2015. Definition of a Novel Pathway Centered on Lysophosphatidic Acid To Recruit Monocytes during the Resolution Phase of Tissue Inflammation. J. Immunol. 195, 1500733. doi:10.4049/jimmunol.1500733

McArthur, S., Juban, G., Gobbetti, T., Desgeorges, T., Theret, M., Gondin, J., Toller-Kawahisa, J.E., Reutelingsperger, C.P., Chazaud, B., Perretti, M., Mounier, R., 2020. Annexin A1 drives macrophage skewing to accelerate muscle regeneration through AMPK activation. J. Clin. Invest. 130, 1156–1167. doi:10.1172/JCI124635

Meng, H., Zhao, H., Cao, X., Hao, J., Zhang, H., Liu, Y., Zhu, M.-S., Fan, L., Weng, L., Qian, L., Wang, X., Xu, Y., 2019. Double-negative T cells remarkably promote neuroinflammation after ischemic stroke. Proc. Natl. Acad. Sci. U. S. A. 116, 5558–5563. doi:10.1073/pnas.1814394116

Misko, T.P., Schilling, R.J., Salvemini, D., Moore, W.M., Currie, M.G., 1993. A Fluorometric Assay for the Measurement of Nitrite in Biological Samples. Anal. Biochem. 214, 11–16. doi:10.1006/abio.1993.1449

Moussa, C., Hebron, M., Huang, X., Ahn, J., Rissman, R.A., Aisen, P.S., Turner, R.S., 2017. Resveratrol regulates neuro-inflammation and induces adaptive immunity in Alzheimer’s disease. J. Neuroinflammation 14, 1. doi:10.1186/s12974-016-0779-0

Nair, S., Sobotka, K.S., Joshi, P., Gressens, P., Fleiss, B., Thornton, C., Mallard, C., Hagberg, H., 2019. Lipopolysaccharide-induced alteration of mitochondrial morphology induces a metabolic shift in microglia modulating the inflammatory response in vitro and in vivo. Glia 67, 1047–1061. doi:10.1002/glia.23587

O’Neill, L.A.J., Pearce, E.J., 2016. Immunometabolism governs dendritic cell and macrophage function. J. Exp. Med. 213, 15–23. doi:10.1084/jem.20151570

Paolicelli, R.C., Bolasco, G., Pagani, F., Maggi, L., Scianni, M., Panzanelli, P., Giustetto, M., Ferreira, T.A., Guiducci, E., Dumas, L., Ragozzino, D., Gross, C.T., 2011. Synaptic pruning by microglia is necessary for normal brain development. Science (80-.). 333, 1456–1458. doi:10.1126/science.1202529

Park, J., Min, J.-S., Kim, B., Chae, U.-B., Yun, J.W., Choi, M.-S., Kong, I.-K., Chang, K.-T., Lee, D.-S., 2015. Mitochondrial ROS govern the LPS-induced pro-inflammatory response in microglia cells by regulating MAPK and NF-κB pathways. Neurosci. Lett. 584, 191–196. doi:10.1016/j.neulet.2014.10.016

Ransohoff, R.M., El Khoury, J., 2015. Microglia in Health and Disease. Cold Spring Harb. Perspect. 8, a020560. doi:10.1016/j.mito.2018.06.006

Ries, M., Loiola, R., Shah, U.N., Gentleman, S.M., Solito, E., Sastre, M., 2016. The anti-inflammatory Annexin A1 induces the clearance and degradation of the amyloid-β peptide. J. Neuroinflammation 1–15. doi:10.1186/s12974-016-0692-6

Salter, M.W., Stevens, B., 2017. Microglia emerge as central players in brain disease. Nat. Med. 23, 1018–1027. doi:10.1038/nm.4397

Scannell, M., Flanagan, M.B., deStefani, A., Wynne, K.J., Cagney, G., Godson, C., Maderna, P., 2007. Annexin-1 and Peptide Derivatives Are Released by Apoptotic Cells and Stimulate Phagocytosis of Apoptotic Neutrophils by Macrophages. J. Immunol. 178, 4595–4605. doi:10.4049/jimmunol.178.7.4595

Schattling, B., Engler, J.B., Volkmann, C., Rothammer, N., Woo, M.S., Petersen, M., Winkler, I., Kaufmann, M., Rosenkranz, S.C., Fejtova, A., Thomas, U., Bose, A., Bauer, S., Träger, S., Miller, K.K., Brück, W., Duncan, K.E., Salinas, G., Soba, P., Gundelfinger, E.D., Merkler, D., Friese, M.A., 2019. Bassoon proteinopathy drives neurodegeneration in multiple sclerosis. Nat. Neurosci. 22, 887–896. doi:10.1038/s41593-019-0385-4

Schloer, S., Hübel, N., Masemann, D., Pajonczyk, D., Brunotte, L., Ehrhardt, C., Brandenburg, L.-O., Ludwig, S., Gerke, V., Rescher, U., 2019. The annexin A1/FPR2 signaling axis expands alveolar macrophages, limits viral replication, and attenuates pathogenesis in the murine influenza A virus infection model. FASEB J. 33, 12188–12199. doi:10.1096/fj.201901265R

Serhan, C.N., 2017. Treating inflammation and infection in the 21st century: new hints from decoding resolution mediators and mechanisms. FASEB J. 31, 1273–1288. doi:10.1096/fj.201601222R

Serhan, C.N., Savill, J., 2005. Resolution of inflammation: the beginning programs the end. Nat. Immunol. 6, 1191–7. doi:10.1038/ni1276

Smith, C.J., Hulme, S., Vail, A., Heal, C., Parry-Jones, A.R., Scarth, S., Hopkins, K., Hoadley, M., Allan, S.M., Rothwell, N.J., Hopkins, S.J., Tyrrell, P.J., 2018. SCIL-STROKE (Subcutaneous Interleukin-1 Receptor Antagonist in Ischemic Stroke): A Randomized Controlled Phase 2 Trial. Stroke 49, 1210–1216. doi:10.1161/STROKEAHA.118.020750

Stempel, H., Jung, M., Pérez-Gómez, A., Leinders-Zufall, T., Zufall, F., Bufe, B., 2016. Strain-specific Loss of Formyl Peptide Receptor 3 in the Murine Vomeronasal and Immune Systems. J. Biol. Chem. 291, 9762–75. doi:10.1074/jbc.M116.714493

Tankou, S.K., Regev, K., Healy, B.C., Cox, L.M., Tjon, E., Kivisakk, P., Vanande, I.P., Cook, S., Gandhi, R., Glanz, B., Stankiewicz, J., Weiner, H.L., 2018. Investigation of probiotics in multiple sclerosis. Mult. Scler. 24, 58–63. doi:10.1177/1352458517737390

Thygesen, C., Larsen, M.R., Finsen, B., 2019. Proteomic signatures of neuroinflammation in Alzheimer’s disease, multiple sclerosis and ischemic stroke. Expert Rev. Proteomics 16, 601–611. doi:10.1080/14789450.2019.1633919

Vital, S.A., Becker, F., Holloway, P.M., Russell, J., Perretti, M., Granger, D.N., Gavins, F.N.E., 2016. Fpr2/ALX Regulates Neutrophil-Platelet Aggregation and Attenuates Cerebral Inflammation: Impact for Therapy in Cardiovascular Disease. Circulation 133, 2169–2179. doi:10.1161/CIRCULATIONAHA.115.020633

Walker, J.M., Kruger, N.J., 2003. The Bradford Method for Protein Quantitation, in: Basic Protein and Peptide Protocols. Humana Press, New Jersey, pp. 9–16. doi:10.1385/0-89603-268-x:9

Wan, M., van der Does, A.M., Tang, X., Lindbom, L., Agerberth, B., Haeggström, J.Z., 2014. Antimicrobial peptide LL-37 promotes bacterial phagocytosis by human macrophages. J. Leukoc. Biol. 95, 971–981. doi:10.1189/jlb.0513304

Weiß, E., Schlatterer, K., Beck, C., Peschel, A., Kretschmer, D., 2019. Formyl-Peptide Receptor Activation Enhances Phagocytosis of Community-Acquired Methicillin-Resistant Staphylococcus aureus. J. Infect. Dis. 221, 668–678. doi:10.1093/infdis/jiz498

Yauger, Y.J., Bermudez, S., Moritz, K.E., Glaser, E., Stoica, B., Byrnes, K.R., 2019. Iron accentuated reactive oxygen species release by NADPH oxidase in activated microglia contributes to oxidative stress in vitro. J. Neuroinflammation 16, 41. doi:10.1186/s12974-019-1430-7

Ye, J., Jiang, Z., Chen, X., Liu, M., Li, J., Liu, N., 2017. The role of autophagy in pro-inflammatory responses of microglia activation via mitochondrial reactive oxygen species in vitro. J. Neurochem. 142, 215–230. doi:10.1111/jnc.14042

Youssef, M., Ibrahim, A., Akashi, K., Hossain, M.S., 2019. PUFA-Plasmalogens Attenuate the LPS-Induced Nitric Oxide Production by Inhibiting the NF-kB, p38 MAPK and JNK Pathways in Microglial Cells. Neuroscience 397, 18–30. doi:10.1016/j.neuroscience.2018.11.030

Yu, Y., Ye, R.D., 2014. Microglial Aβ Receptors in Alzheimer’s Disease. Cell. Mol. Neurobiol. 35, 71–83. doi:10.1007/s10571-014-0101-6

Zabala, A., Vazquez-Villoldo, N., Rissiek, B., Gejo, J., Martin, A., Palomino, A., Perez-Samartín, A., Pulagam, K.R., Lukowiak, M., Capetillo-Zarate, E., Llop, J., Magnus, T., Koch-Nolte, F., Rassendren, F., Matute, C., Domercq, M., 2018. P2×4 receptor controls microglia activation and favors remyelination in autoimmune encephalitis. EMBO Mol. Med. 10. doi:10.15252/emmm.201708743

Zhang, J., Malik, A., Choi, H.B., Ko, R.W.Y., Dissing-Olesen, L., MacVicar, B.A., 2014. Microglial CR3 activation triggers long-term synaptic depression in the hippocampus via NADPH oxidase. Neuron 82, 195–207. doi:10.1016/j.neuron.2014.01.043

Zhou, L.J., Peng, J., Xu, Y.N., Zeng, W.J., Zhang, J., Wei, X., Mai, C.L., Lin, Z.J., Liu, Y., Murugan, M., Eyo, U.B., Umpierre, A.D., Xin, W.J., Chen, T., Li, M., Wang, H., Richardson, J.R., Tan, Z., Liu, X.G., Wu, L.J., 2019. Microglia Are Indispensable for Synaptic Plasticity in the Spinal Dorsal Horn and Chronic Pain. Cell Rep. 27, 3844-3859.e6. doi:10.1016/j.celrep.2019.05.087

